# Inflammatory responses is induced in pulmonary embolic lung: evaluation by a reproducible pulmonary embolism model in mice

**DOI:** 10.1101/2022.07.29.501960

**Authors:** Honoka Okabe, Haruka Kato, Momoka Yoshida, Mayu Kotake, Ruriko Tanabe, Yasuki Matano, Masaki Yoshida, Shintaro Nomura, Atsushi Yamashita, Nobuo Nagai

**Affiliations:** Laboratory of Animal Physiology, Division of Animal Bioscience, Faculty of Bioscience, Nagahama Institute of Bio-Science and Technology; Division of Animal Bioscience, Faculty of Bioscience, Nagahama Institute of Bio-Science and Technology; Skin Physiology Laboratory, School of Bioscience and Biotechnology, Tokyo University of Technology; Department of Pathology, Miyazaki University School of Medicine

**Author notes:** Corresponding Author: Nobuo Nagai, PhD. Laboratory of Animal Physiology, Division of Animal Bioscience, Faculty of Bioscience, Nagahama Institute of Bio-Science and Technology.

**Keywords:** Pulmonary embolism, inflammation, angiography, IL-6, macrophage, TNF-α

## Abstract

**Background:** Since previously established models of pulmonary embolism showed a large variability in the degree of ischemia, it is difficult to assess the pathophysiological response in the lung after embolization. Here, we established a model of pulmonary embolism by certain amount of relatively small thrombi, in which the degree of ischemia was reproducible.

**Methods:** Thrombi with a maximum diameter of 100 μm or 500 μm were administered intravenously under anesthesia, and the survival ratio at 4 hours was evaluated. The location of thrombi in the lung was visualized by administration of fluorescent-labeled thrombus, and the hemodynamics of the lung after administration of thrombi was evaluated. CT angiography was also performed to evaluate the ratio of the embolized vessels. In addition, cytokine mRNAs was quantified 4 hours in embolized lung. Immunohistochemical analysis for interleukin (IL)-6 and CD68 as a marker of macrophages were also performed.

**Results:** It was found that mice with 100 μm clots, but not with 500 μm clots, showed a dose-dependence of survival between 2.3 μL/g and 3.0 μL/g at 4 hours from embolization induction. In mice treated with 2.5 μL/g of 100 μm thrombus, thrombi were located in the peripheral region of the lung, which was consistent with the disruption of blood circulation the peripheral region. In addition, about 60% of the vessels with a diameter of less than 100 μm were occluded in these mice. In the lungs after 4 hours of embolization, IL-6 mRNA and tumor necrosis factor (TNF)-α mRNA were significantly higher and lower than control lungs. IL-6 was expressed in CD68-positive macrophages in both embolized and control lungs after 4 hours of embolization, and the number of each positive cells were comparable in both embolized and control lungs.

**Conclusions:** These results show that the pulmonary embolization model induced by a certain amount of small thrombus is useful for evaluating the pathological responses in the embolized lung. Furthermore, it was found that IL-6 expression was increased in macrophages in the embolized lung, indicating that inflammatory responses may contribute to the pathogenesis of pulmonary embolism.

## Backgrounds

Pulmonary embolism is a disease in which a thrombus formed in the deep veins reaches the pulmonary artery and becomes an embolus. For this study, various models have been established by exogenous thrombus injection or endogenous thrombus induction by administration of coagulation factors such as thrombin or tissue factor (1–5). Although these models are useful for evaluating the effect of thrombolytic agents and endogenous fibrinolytic activity in dissolving pulmonary emboli (3–5), they are not suitable for assessing the pathogenesis of pulmonary emboli because of the large variability in the pathogenesis associated with pulmonary emboli.

In this report, we established a model that induces the pathogenesis of pulmonary embolism in a thrombus dose-dependent manner by inducing many relatively small pulmonary artery emboli with a fixed dose of small thrombus.

First, we studied the effect of clot size on pathological responses in pulmonary embolic model, and found that the survival rate at 4 hours after administration was dose-independent with clot-diameter of up to 500 μm, whereas it was dose-dependent with clot-diameter of up to 100 μm in the range from 2.0 μL/kg to 3.0 μL/kg.

Then we investigated the hemodynamics and the induction of proinflammatory cytokines in the lung after pulmonary embolization by clots with a diameter of up to 100 μm.

## Materials and Methods

### Animals

All animal experimental procedures were approved by the Committee on Animal Care and Use of the Nagahama Institute of Bio-Science and Technology (Permit Number: 017). The animal studies were performed in accordance with institutional and national guidelines and regulations, and the ARRIVE (Animal Research: Reporting of In Vivo Experiments) guidelines (https://www.nc3rs.org.uk/arrive-guidelines). In whole experiments, 15 C57BL/6J male mice (CLEA Japan, Tokyo, Japan) 12-14 weeks old and weighing 25-30 g were used for plasma sampling. Fifty-eight BALB/c female mice (CLEA Japan), 12-16 weeks old and weighing 20-28 g, were used for pulmonary embolism induction. In these were used for analysis as follows; 26 for clot dose response, 3 for clot distribution with fluorescence-labeled clots, 3 for blood perfusion in embolized lung, 14 for computer tomography (CT) angiography, and 12 for mRNA measurement.

### Clot formation

Clot was made from mouse plasma obtained from C57Bl/6J male mice. Under anesthetization by a combination anesthetic prepared with 0.3 mg/kg of medetomidine, 4.0 mg/kg of midazolam, and 5.0 mg/kg of butorphanol, blood taken from hart was mixed with 10% sodium citrate for anticoagulation. After centrifugation, plasma from 15 mice were mixed and stored at −80 °C until use. The concentration of fibrinogen was 270 mg/dL, which was measured by a commercial kit for human (AssayPro, MO, USA). For clot formation, 100 μL of plasm was mixed with 2.5 μl of 1M CaCl_2_ (nacalai tesqu, Kyoto, Japan) and 10 μl of human thrombin (nacalai tesqu), and incubated at 37 °C over 1 hour. Then 400 μl of 4% sodium citrate was added for preventing further coagulation. The clot was dissected by small scissors and crashed by a sonicator (UD-201, Tomy, Tokyou, Japan) over certain seconds. The size was checked under microscope. For visualizing the clots, 0.67mg/mL of human Fbg-TM488 (Invitogen, MA, USA) was added when clot was made.

### Pulmonary embolism

Pulmonary embolism was induced by injection of clots in BALB/c mice. Mouse was anesthetized by isoflurane and kept on a heat pad at 37 °C. The skin of neck was incised and a catheter filled with clot solution was inserted to Juglar vein. A certain amount of clot solution was infused through the catheter over around 10 seconds. Then the catheter was withdrawn and skin was replaced and sutured. For evaluating the survival ratio, animals were kept under isoflurane until 4 hours. If breath stopped more than 30 second, we decided the animal die and euthanized by intraperitoneal injection of 200 mg/kg of sodium secobarbital.

### Hemodynamics

To evaluate the perfusion region of the lung after clot injection, mouse was perfused with Evans blue. Under isoflurane anesthesia, left atrium was incised and 10mL of 4% Evans Blue in saline was perfused from left ventricle of the heart in mice. Then, lung vessels were ligated, the lung was taken, and photographed.

### CT imaging

To assess the vessel occlusion, we performed CT angiography after clot injection. Under isoflurane anesthesia, left atrium was incised and 10mL of contrast media (ref) was perfused from left ventricle of the heart in mice. Then, lung vessels were ligated, the lung was taken, CT imaging was performed by mCT_2 (Rigaku, Tokyo, Japan). Parameters used for the CT scans were as follows: tube voltage: 90 kV; tube current: 200 μA; axial field of view (FOV): 30 mm, with an inplane spatial resolution of 10 μm. Total scan time was approximately 3 min. The image data were stored in DICOM format. Then, the numbers of brunches were counted using AIVIA analysis system (DRVISION, Bellevue, USA) performed with ORIENT SYSTEM, INC. (Tokyo, Japan). In this analysis, three dimensional image was reconstructed in each animal and all branches in the image were divided in two groups by their diameters; over or under 100 μm. Then the number of brunches with diameter less than 100 μm were counted.

### Quantitative PCR

Expression of mRNA of cytokines were measured. Four hours after pulmonary embolism, mice were euthanized by intraperitoneal injection of 200 mg/kg of sodium secobarbital. The lung was taken, frozen by liquid nitrogen, and stored at −80 °C until measurement. The lung was mixed with trizol (nacalai tesque) and homogenized (beads homogenizer). Total RNA was isolated by using a RNeasy Micro Kit (Qiagen, Hilden, Germany) according to the manufacturer’s instructions. Reverse transcription was performed using a cDNA Reverse Transcription Kit (Toyobo, Tokyo, Japan). Quantitative real-time PCR was performed by TB Green Fast qPCR Mix (Takara Bio Inc. Kusatsu, Japan) on a PCR System (Roche, Basel, Switzerland). The PCR primers are listed in Table 1. The specific mRNA amplification of the target was determined as the Ct value followed by normalisation with the GAPDH mRNA level. The value was represented by the ratio against that in control lung.

**Table 1.**
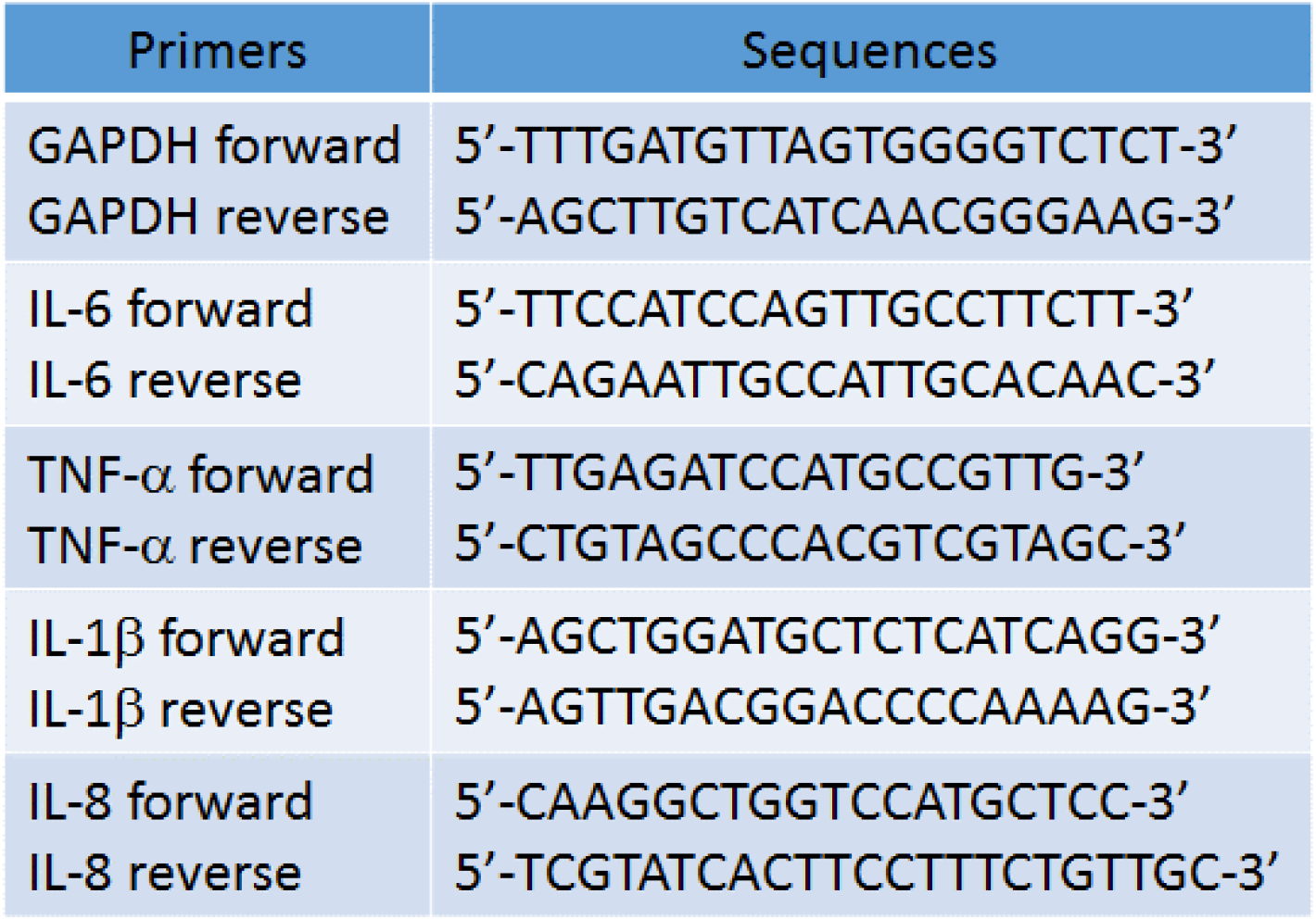
PCR primers

### Immunohistochemistry

Four hour after pulmonary embolism induction, mice were perfused with 4 % paraformaldehyde under sebobarbital anesthesia. Their lungs were removed, embedded in paraffin, and sliced into 5mm thickness sections, which were then immunostained as described elsewhere. Briefly, a section and the adjacent section were treated with anti-IL-6 and anti-CD68 as a marker of macrophages, respectively. After treatment with an appropriate secondary antibody conjugated with peroxidase, the immunoreactivity was visualized by diaminobenzidine colouration. Then, sections were counterstained by hematoxylin. Microphotographs were taken under a microscope.

### Statistics

Statistical analysis was performed by Student’s t-test. P-values less than 0.05 were considered significant.

## Results

### Effect of clot size and clot dose on survival ratio

First, we studied the clot-dose response on survival ratio in by using clots in two different size. When the clots in which the maximum size was around 500 μm were used, the survival ratio was not associated with its dose. When the clots in which the maximum size was around 100 μm were used, the survival ratio was associated with its dose, that is, survival ratio increased dose response manner, from 2.3 μL/g to 3.0 μL/g (Table 2). Based on these results, we chose the clots in which the maximum size was around 100 μm for further experiments.

**Table 2.** Dose response in different clot size

| Clot size 500µm |  |  |  | Clot size 100µm |  |  |  |
| --- | --- | --- | --- | --- | --- | --- | --- |
| Dose (µL/g) | Live | Dead | % Live | Dose (µL/g) | Live | Dead | % Live |
| 2.0 | 2 | 2 | 50 | 2.3 | 3 | 0 | 100 |
| 2.3 | 1 | 3 | 25 | 2.5 | 3 | 1 | 75 |
| 3.0 | 2 | 2 | 50 | 2.8 | 1 | 2 | 33 |
|  |  |  |  | 3.0 | 1 | 3 | 25 |

### Clot distribution in embolized lung

In embolized lung with 2.5 μL/g clot solution mixed with human Fbg-TM488, several fluorescent signals were observed. These signals were distributed relatively near the edge of lung leafs (Figure 1).

**Figure 1.**
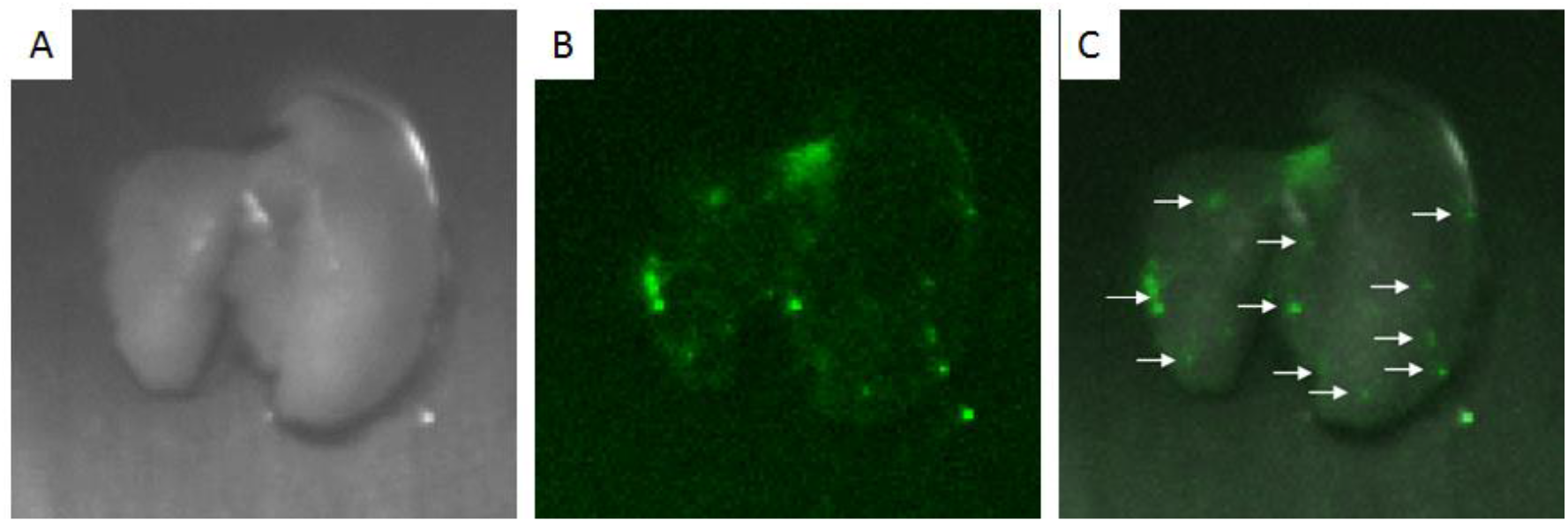
Distribution of embolic clots in the lung. Reflection image (A) and fluorescence image (B) and their marge image (C) of clot-injected lung. Arrows indicate clot distributed near the edge of each leaves.

### Perfused region in embolized lungs

In normal lung, whole tissue was stained with Evans blue (Figure 2A), indicating that blood perfusion was kept normally in all area of the lung. In lung with pulmonary embolization of 2.5 μL/g clot solution, however, the edge of major lung leafs with whole area of right sub leaf were not perfused (Figure 2B). We found similar results in other two animals in each group (data not shown).

**Figure 2.**
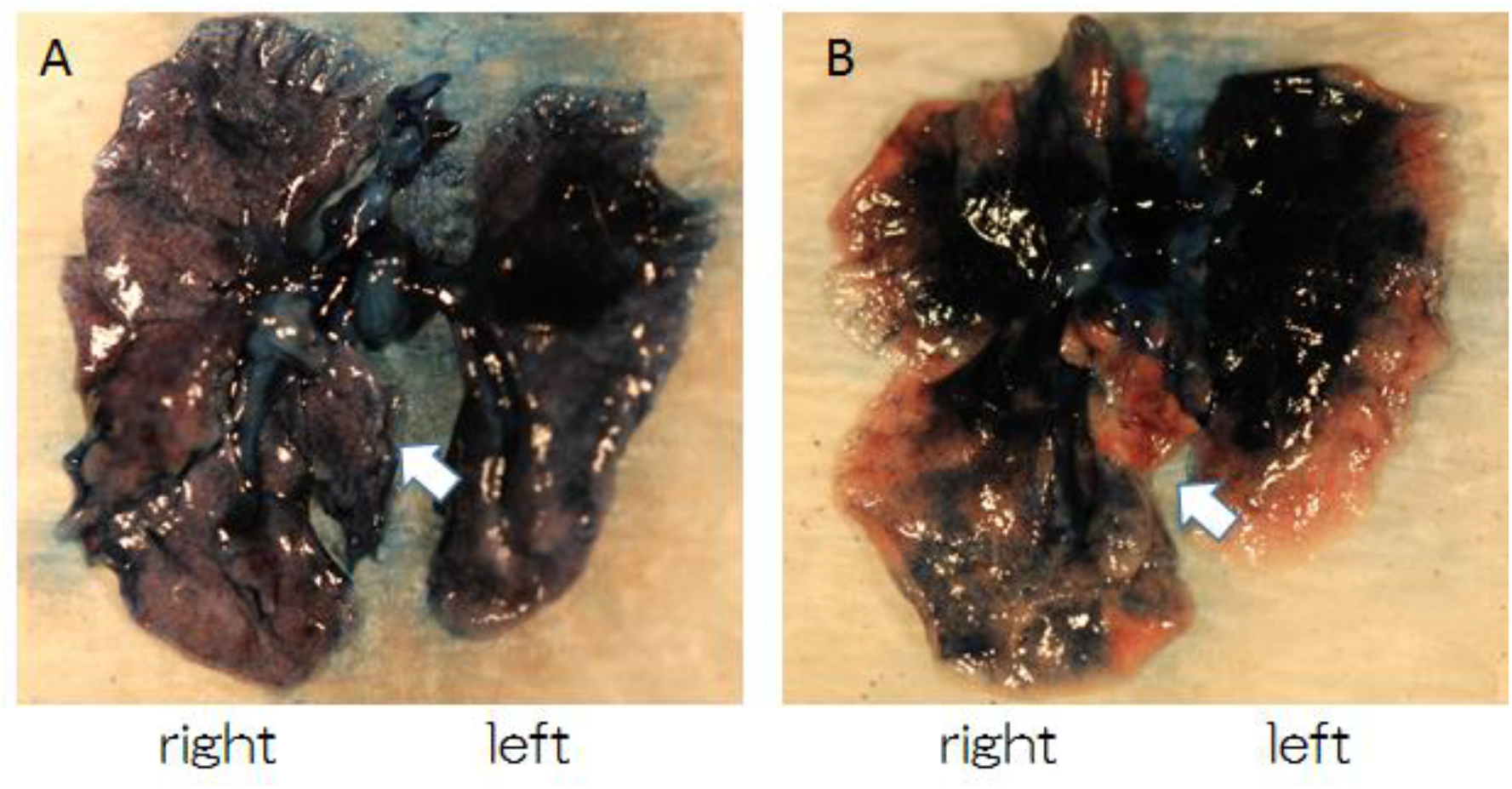
Perfusion in the lung Photographs of lungs without clot-injection (A) and with clot-injection (B) after perfusion of Evans blue from right ventricle to left atrium. Arrows indicate right sub leaf. Non perfusion area was distributed widely at the edge of all leafs of lung in clot-injected mice.

### CT angiography

In normal lung, we could visualized vessels of which the diameter was around 50 μm by using the titanium contract (Figure 3A). In embolized lung with 2.5 μL/g clot solution, the number of vessels decreased (Figure 3B). In addition, the vessel branch of right sub leaf was shortened than that in normal lung (Figure 3A and B). The numbers of vessel branch with diameter less than 100 μm was significantly less in embolized lung than normal lung.

**Figure 3.**
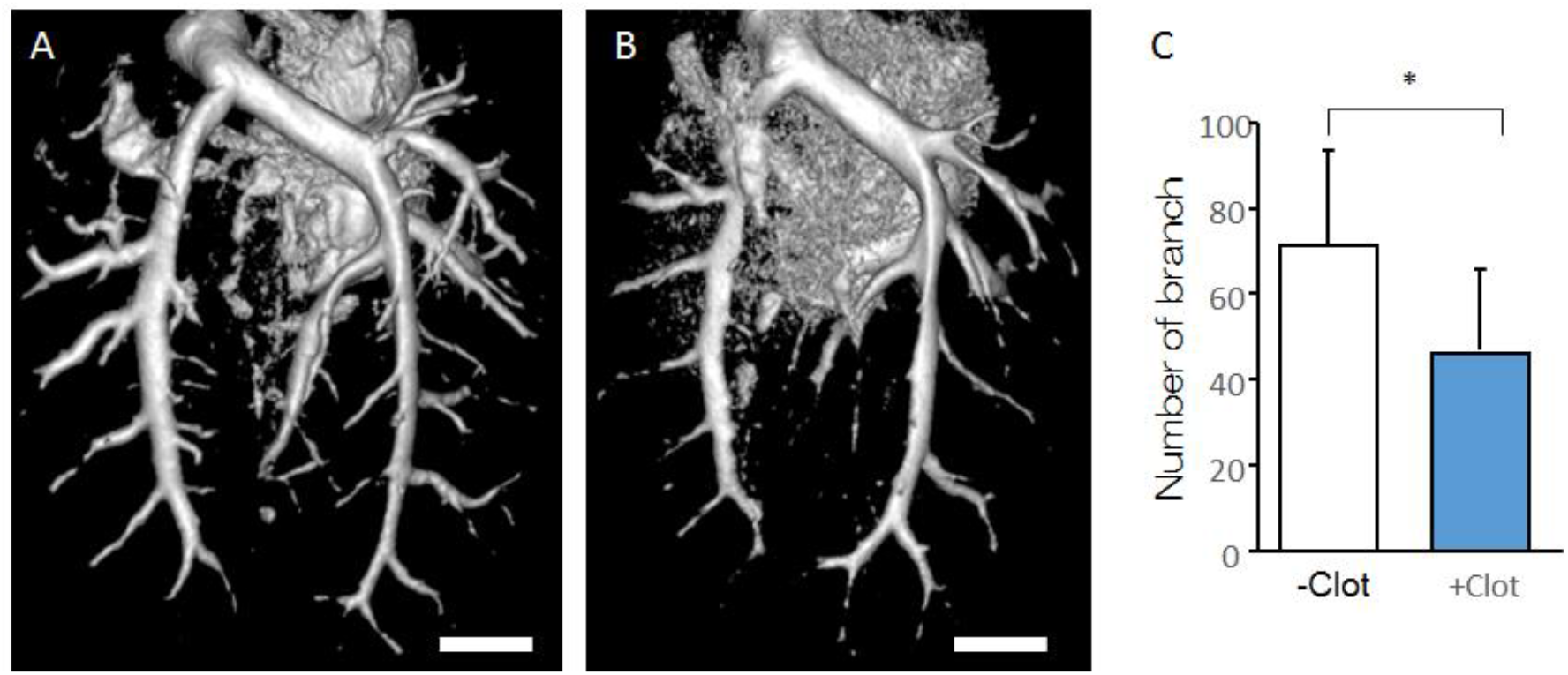
CT angiography in the lung CT angiography of lungs from mouse without clot injection (A) and with clot injection (B). The number of brunch with diameter less than 100 μm (C). Each group consists of 7 mice, respectively. Data represent mean and SD. *:*p*<0.05.

### mRNA expression of cytokines

To evaluate the effect of embolism on gene expression relating hypoxia response and cytokines, we measured mRNA of IL-1β, IL-6, IL-8 and TNF-α (Figure 4). At 4 hours after clot injection, the amount of mRNA of IL-1 β and IL-8 were comparable in both normal lung and enbolized lung, respectively. In addition IL-6 mRNA increased and TNF-α mRNA decreased at 4h after clot injection, which were both significantly.

**Figure 4.**
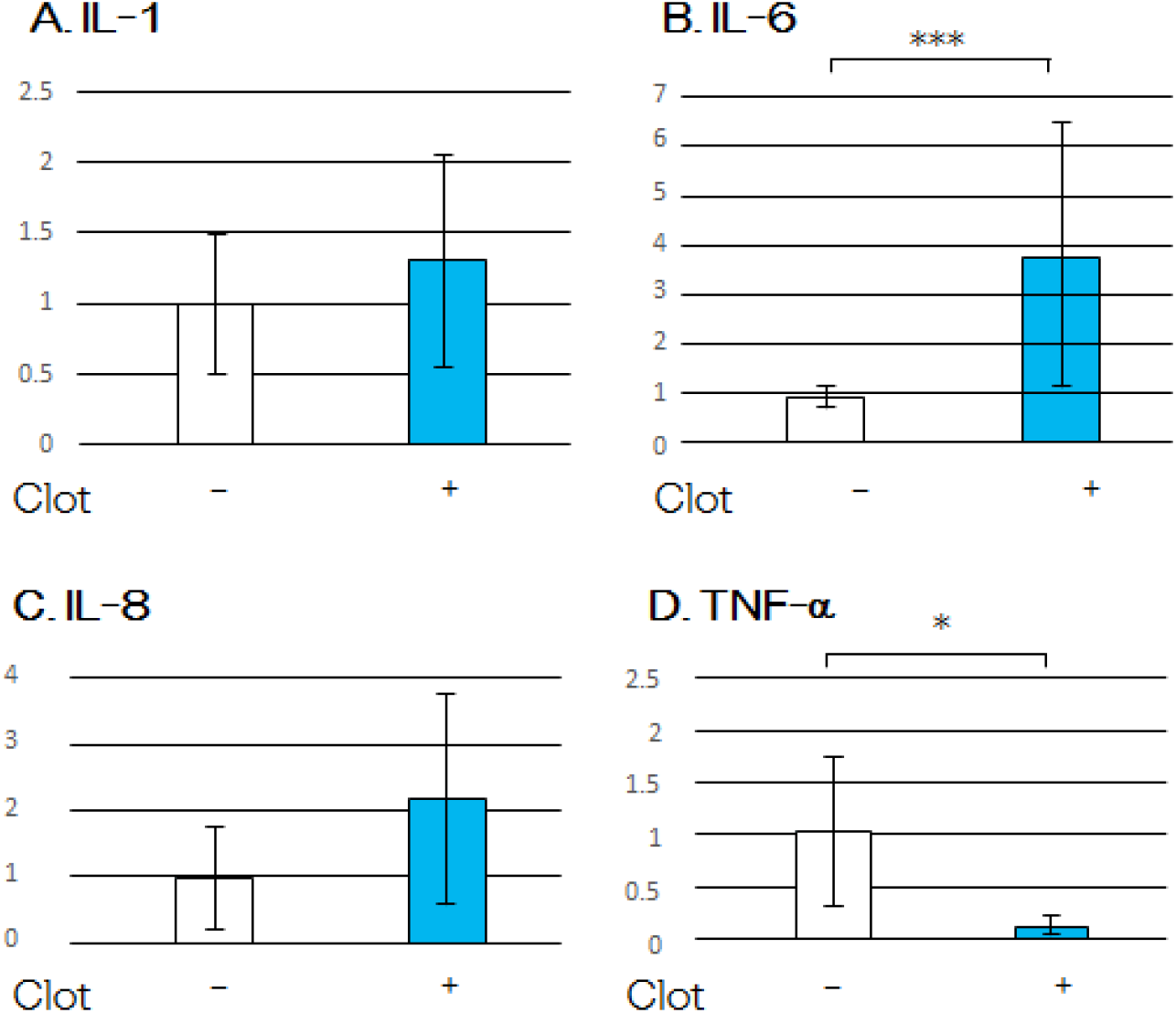
Effect of clot injection on mRNA expression of HIF-1a and cytokines. mRNA levels of IL 1β (A), IL-6 (B), IL-8 (C) and TNF-α (D) at 4 hours after clot injection. Data represent mean and SD of relative value against mice without clot injection (-Clot). Each group consists of 6 mice, respectively. *:*p*<0.05, ***:*p*<0.001.

### Immunohistochemistry of IL-6 and CD68

On immunohistochemical analysis for IL-6, positive cells were observed in both control lung and embolized lung (Figure 5A and B). On immunohistochemical analysis for CD68, a marker of macrophage, positive cells were distributed consistently with IL-6 immunostained cells in adjacent sections (Figure 5C and D). The numbers of both IL-6 immunostained cell (Figure 5E) and CD68 immunostained cells (Figure 5F) were comparable between the control lung and the embolized lung. In addition, the numbers of IL-6 immunostained cell and CD68 immunostained cell are close in the both control lung and the embolized lung.

**Figure 5.**
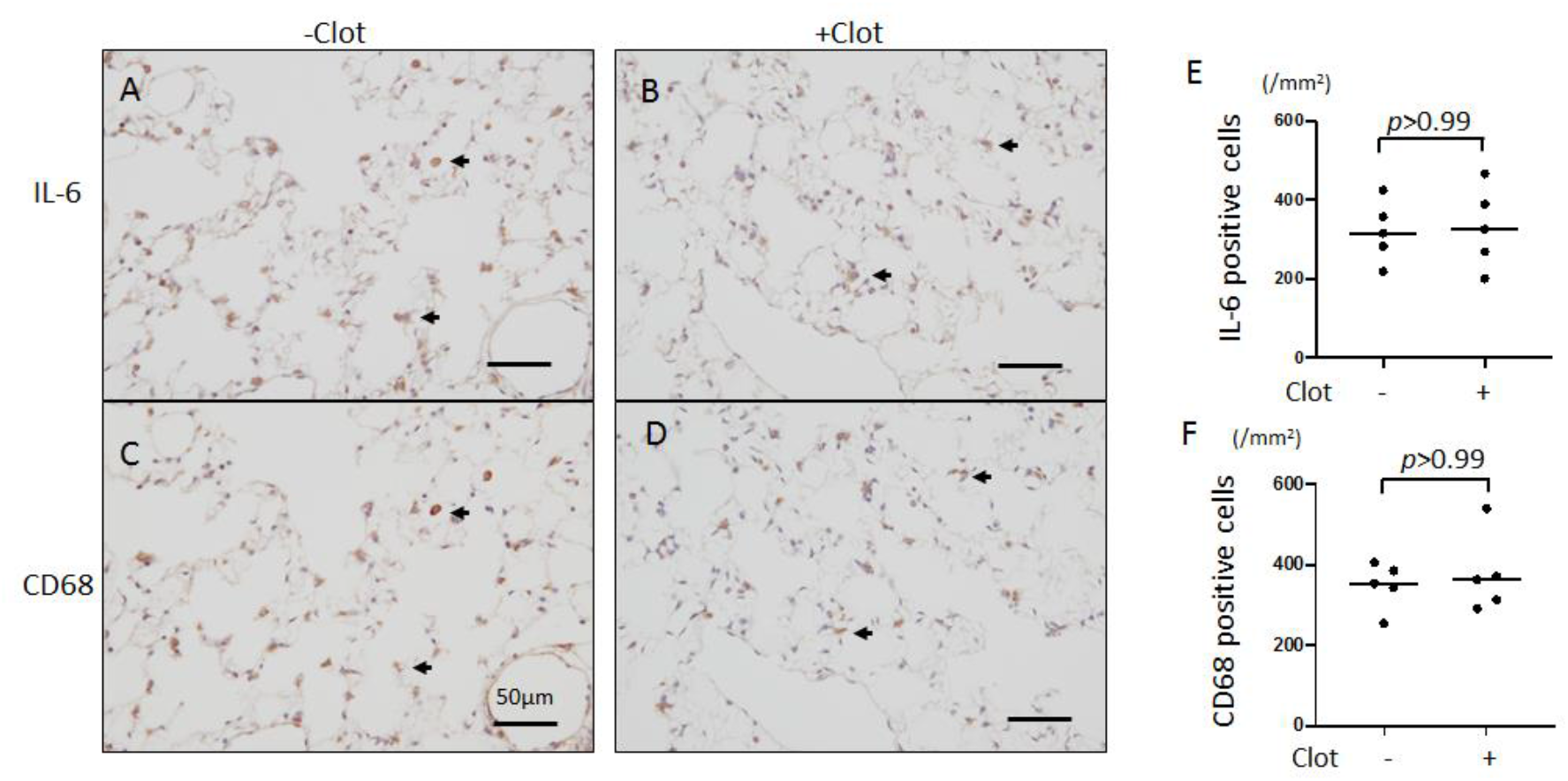
Distribution of immunoreactive cells against IL-6 and CD68. Photographs of lung sections stained with anti-IL-6 or anti-CD68 in adjacent sections in mice without (-Clot) or with (+Clot) clot injection (A). Arrows indicate double positive cells against both IL-6 and CD68. Most of positive cells ware overlapped. The numbers of positive cells in each staining in individual mice were shown in B. Each dot indicate the number of positive cells/mm^2^ of sections in each animal.

## Discussion

In this study, we established a pulmonary embolus model by administering a fixed amount of small clots, in which the survival rate increased in a clot dose-dependent manner. Therefore, we succeeded to maintain the animals with pulmonary embolism to be alive with almost maximum ischemic damage, measure the expression cytokine mRNA. These results indicate the usefulness of this model for analyzing the pathological responses after pulmonary embolism.

Several pulmonary embolization models have been established, in which embolization was induced by several large clots or generated clots in the body (1–5). These models are useful for evaluating the effects of thrombolytic agents and endogenous fibrinolytic activity in dissolving pulmonary emboli. Since their ischemic tendency varied greatly, however, they were not useful for analysis of the pathological responses after pulmonary embolism. In the present study, we also found that there was no correlation between survival rate at 4 hours and dose in pulmonary embolus caused by a relatively large thrombus with a maximum diameter of 500 μm. On the other hand, for pulmonary embolus caused by a thrombus with a maximum diameter of 100 μm, the survival rate decreased in a dosedependent manner between 2.0 μL/g and 3.0 μL/g. This result suggests that, for the purpose of investigating the quantitative effects of pulmonary embolization on pulmonary occlusion, a thrombus with a maximum diameter of 100 m was suggested to be useful.

Since oxygen and nutrients are supplied to the lung tissue via the bronchial arteries, the obstruction of the pulmonary arteries does not affect the blood supply to the lungs. Therefore, the effects of ischemia in the lung by pulmonary emboli are expected to depend not on the location of the pulmonary embolus but on the size of the occluded area of the lung. In addition, since the downstream area of the large branches of the pulmonary artery varies from branch to branch, the size of the occluded area of the lung will also vary depending on the branch being occluded by several large thrombi, and therefore, survival ratio will be independent of the amount of thrombus. On the other hand, in the case of pulmonary embolism caused by many small thrombi, the number of occluded small branches would increase depending on the amount of thrombus and result in an increase in the occlusion volume and a decrease in the survival rate. To characterize the pulmonary embolization by a thrombus with a maximum diameter in 100 μm, the distribution of thrombus, blood perfusion area in lung, and CT angiography were performed. It was confirmed that several thrombi were localized in the marginal region of lung lobes. Blood perfusion in the lung lobes o was also inhibited in the marginal region, which were consistent with the distribution of thrombi. Furthermore, the number of vessel branches below 100 μm was significantly decreased in embolized lung to about 60% of that in control lung when 2.5 μL/g of thrombus was administered. Considering that the survival rate with 2.5 μL/g of thrombus was 75%, it is suggested that the survival rate decreases when the number of vessel branches below 100 μm falls below 60%.

It was found that the expression levels of both IL-1β and IL-8 mRNA were comparative in both the lung 4 hours after administration of 2.5 μL/g thrombus and control lung, whereas IL-6 mRNA was significantly increased. IL-6 is a cytokine that plays an important role in inflammatory reactions, which induces chemokine production in various cells, and enhances the expression of adhesion molecules in vascular endothelial cells. In the lung, it has been shown that IL-6 increased significantly during the acute inflammatory response including bronchoscopic alveolar lavage (6) and acute pneumonia by lipopolysaccharide within 3 hours (7), which were thought to deteriorate the damage. In addition, it has been reported that ischemia also increases IL-6 in the brain, heart and kindney in the early stages (8–10) indicating that IL-6 contributes to the acute inflammatory response associated with ischemia. The current findings that IL-6 mRNA was upregulated in the embolized lung, therefore, also suggest that IL-6 is expressed in the embolized lung may be involved in the pathogenic responses in acute phage of pulmonary embolism. Immunohistological studies revealed that macrophages expressed IL-6, which was increased in the lung 4 hours after pulmonary embolization. Expression of IL-6 in alveolar macrophages has also been reported systemic sclerosis (11) and pneumonia (12). In contrast, the number of both IL-6 immunoreactive cells and macrophages in the lung was similar to that in the control lung at 4 hours after pulmonary embolization. These results suggest that the increase in IL-6 mRNA may be due to an increase in each macrophage. The function of IL-6 in pathogenesis varies by organ and disease. IL-6 exacerbates the pathology in the acute phase of cerebral infarction (13), whereas it plays a protective role in renal ischemia-reperfusion injury (14). The contribution of IL-6 in the pathogenesis of pulmonary embolism is a subject for further investigation.

We also found that TNF-α mRNA level was significantly lower in embolized lungs than in control lungs. TNF-α is mainly expressed in macrophage, which is upregulated by hypoxia (15, 16). Since it has been reported that human alveolar macrophages have a higher TNF-a production capacity than peripheral blood monocytes (17), however, alveolar macrophages are thought to express relatively high levels of TNF-α even under normal condition. The decrease in TNF-α in embolized lung may be due to effects by pulmonary embolization other than hypoxia, which may have affected alveolar macrophage function and reduced TNF-α expression. The pathophysiological roles of the decrease in TNF-alpha expression in acute phase of embolized lungs is a subject for further investigation.

## Conclusions

We established a novel murine pulmonary embolism model by relatively small clots, in which ischemic damage is reproducible and the survival ratio showed a clot dose-dependent manner. By this model, it was shown that inflammatory responses associating with increase in IL-6 mRNA expression and decrease in TNF-α mRNA expression was induced in embolized lung at 4 hours.

## List of Abbreviations

IL: interleukin
TNF: tumor necrotic factor
CT: Computer Tomography
FOV: field of view

## Declarations

Ethics approval: All animal experimental procedures were approved by the Committee on Animal Care and Use of the Nagahama Institute of Bio-Science and Technology (Permit Number: 017). The animal studies were performed in accordance with institutional and national guidelines and regulations, and the ARRIVE (Animal Research: Reporting of In Vivo Experiments) guidelines (https://www.nc3rs.org.uk/arrive-guidelines).

- Consent for publication: no indivdial person’s data is included
- Availability of data and materials: The datasets used and/or analysed during the current study are available from the corresponding author on reasonable request.
- Competing interests: we have no conflict of interest.
- Funding: This study was supported by a Grant-in-Aid for Scientific Research (C) (grant 15K08194), MEXT-Supported Program for the Strategic Research Foundation at Private Universities (grant 1201037) by the Ministry of Education, Culture, Sports, Science and Technology, and The Research Promotion Grant of The Japanese Society on Thrombosis and Hemostasis.
- Authors’ contributions are as follows: Honoka Okabe, Haruka Kato Momoka Yoshida and Yasuki Matano: maintain and experiments in animals, Ruriko Tanabe and Shintaro Nomura: CT angiography study, Masaki Yoshida: qPCR analysis, Atsushi Yamashita: Immunohistochemical analysis, Nobuo Nagai: conduct of all experiments.

